# Differences in Ion-RNA binding modes due to charge density variations explain the stability of RNA in monovalent salts

**DOI:** 10.1101/2022.02.12.480180

**Authors:** Anja Henning-Knechtel, Devarajan Thirumalai, Serdal Kirmizialtin

## Abstract

The stability of RNA increases as the charge density of the alkali metal cations increases. The molecular mechanism for the stability does not exist. To fill this gap, we performed all-atom molecular dynamics (MD) pulling simulations to dissect the microscopic origin of this phenomenon. We first established that the free energy landscape obtained in the simulations is in excellent agreement with the single-molecule optical tweezer experiments. The origin of the stronger stability in *Na*^+^ compared to *K*^+^ is found to be due to the differences in the charge-density related binding modes. The smaller hydrated *Na*^+^ ion preferentially binds to the highly charged phosphates. In contrast, the larger *K*^+^ ions interact with the major grooves. As a result, the electrostatic repulsion between the phosphate groups is reduced more effectively by *Na*^+^ ions. Because the proposed mechanism is generic, we predict that the same conclusions are valid for divalent alkaline earth metal cations.

## Introduction

RNA folding is a complex process that is determined not only by the nucleotide sequence but also is driven by cations that are needed to soften the electrostatic repulsive interactions between the negatively charged phosphate groups.^1,2^ Although there are exceptions,^3–5^ in general, independently stable secondary structures form first, followed by consolidation of tertiary interactions that stabilize the folded state. Several studies^6–12^ have shown that RNA undergoes dramatic compaction during folding, as the salt concentration is increased. The entropy loss due to secondary structure formation is primarily compensated by the gain in hydrogen bonding and base stacking interactions involving the nucleotides. Condensation of cations onto the phosphate groups is required for RNA to fold.^1,13–17^ Thus, the formation of a stable folded RNA involves an interplay of a number of factors (RNA conformations, size, shape, and valence of cations, and hydration effects), which makes it difficult to decipher the principles governing the folding of RNA.^2^

Analyses of RNA structures and simulations have shown that cations bind to discrete sites on RNA, rather than non-specifically.^18–24^ Optical melting experiments, small angle X-Ray scattering, NMR and gel electrophoretic studies suggest that there is a complicated interplay between RNA conformations and charge density.^25–33^ In addition, single molecule optical and magnetic tweezers experiments^34–38^ and atomic force microscopy^39^ reported a cation-size dependent stabilization of RNA hairpins.^34^ These experiments show higher RNA stability in NaCl solution than in KCl solution. Experimental and theoretical studies suggest that the smaller hydrated monovalent cations are more effective in RNA stabilization. A conclusion of general validity is that cation charge density is a dominant factor in affecting RNA stability.^26,27^

Various mechanisms have been proposed to explain the charge density induced stability in RNA folding.^10,16,26,40–42^ If RNA is more flexible in KCl than in NaCl, the entropic stabilization of the unfolded state would result in a smaller free energy difference between the native and unfolded states in *K*^+^ ion. Another possibility is that the smaller size of *Na*^+^ ions leads to stronger binding to the RNA surface, which could induce further compaction of RNA, which would be consistent with SAXS experiments on a ribozyme.^10^ However, the molecular basis of the postulated charge density, *ρ_c_*, hypothesis remains unclear. Because the results of a large number of experiments on RNA stability hinges on the use of *ρ_c_* as the primary factor in determining RNA stability, providing a molecular explanation would greatly advance our understanding of ion-RNA interactions. Doing so would require atomically detailed simulations that quantitatively accounts for the coupling between RNA conformations and fluctuations in the fields created by the cations.

Here, we use all-atom simulations in explicit water to investigate the impact of Na^+^ and K^+^, with drastically different *ρ_c_* values, on RNA folding. For illustration purposes, we consider the mechanical stability of HIV-1 TAR, a ubiquitous RNA that forms a helixjunction-helix structure. Using mechanical force (*f*) to unfold the hairpin, as was done in single molecule pulling experiments,^34^ we monitor the interactions between the cation and RNA as it unfolds. We first show that the simulations capture the results obtained in experiments,^34^ which showed that HIV-1 TAR is more stable in *Na*^+^ than in *K*^+^. We then determined the microscopic basis of the impact on the stability of HIV-1 TAR in the presence of the two cations. There are marked differences between the binding preferences of the cations to the RNA surface depending on their size, and hence *ρ_c_*. Potassium ions predominantly bind to the major grooves with high affinity. In sharp contrast, the smaller sodium ions, with higher charge density, are localized on the phosphate backbone. These differences in the preferential binding resulted in different coordination numbers. Sodium ions, which are exposed to a larger surface, have a higher number of cation coordination, and hence results in higher thermodynamic stability of the RNA. A broader principle that emerges from this study is that for given ion valence and ion shape, the cation with the highest charge density can bind most tightly to the negatively charge phosphate groups and therefore impart the greatest stability on the folded RNA.

## Results and Discussion

To investigate the energy landscape of RNA subject to mechanical force, and to study the role of monovalent cation size in RNA stabilization, we chose the HIV-1 TAR RNA hairpin, which was the subject of investigation in the pulling experiments by Vieregg, et. al.^34,36^ In Fig. 1A-B we show the sequence and structure of the 52-nt long HIV-1 TAR RNA. The structure consists of a 26-nt extended stem (gray), connected to a 4-nt lower stem (red) with a 3-nt apical loop (green) followed by a 4-bp upper stem (yellow) that is bridged by a 6-nt bulge region (purple). The simulation box with the direction of the pulling coordinate is shown in Fig. 1C. This construct enabled us to directly compare our findings to mechanical pulling experiments. Direct comparison of available experimental data allowed to asses the validity of the theoretical findings.

**Figure 1:**
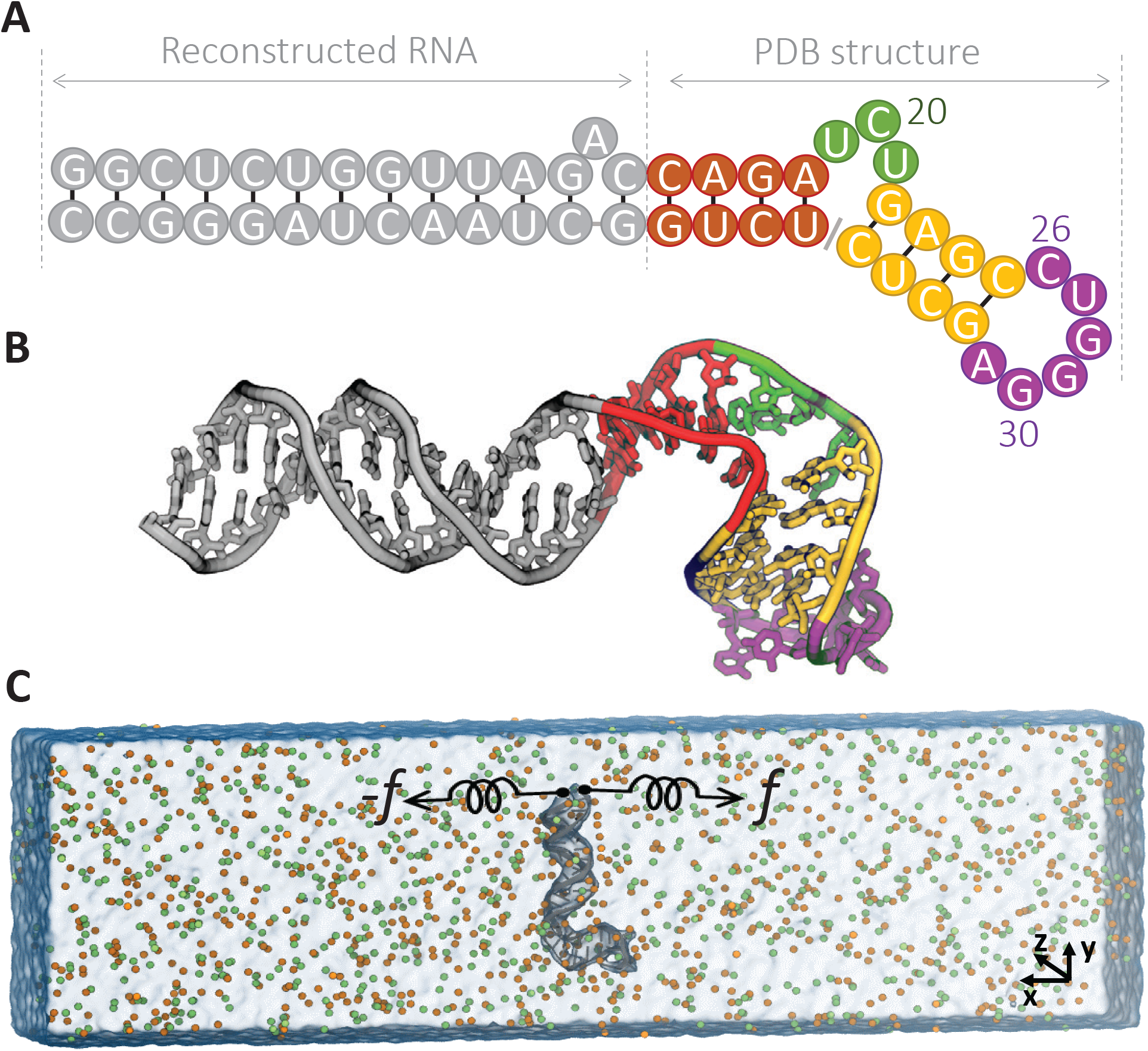
Schematic of molecular simulation set-up for HIV-1 TAR hairpin. A) RNA sequence. B) The structure of the HIV-1 TAR hairpin under study consists of an extended stem (gray), connected to lower stem (red) with apical loop (green) followed by upper stem (yellow) that is bridged by bulge region (purple). C) Set-up for the MD simulation in 400 mM KCl, *K*^+^ (orange) and *Cl*^−^ (green). The RNA hairpin (grey) is aligned perpendicular to the *x*-axis and the termini nucleotide are pulled.

### Cation-dependent energy landscapes

We constructed the energy landscape along the mechanical pulling coordinate using the umbrella sampling approach (detailed in the *Computational Methods* section of the Supporting Information).^43^ The effect of applied force, *f*, on the free energy, projected onto the molecular extension *F_f_* (*x*) is calculated using, *F_f_* (*x*) = *F*_0_(*x*) – *fx*, where *F*_0_(*x*) is the free energy at zero force. Our results, plotted in Fig. 2A-B, display a staircase-like energy landscape, where the intermediate states become visible only if *f* ≠ 0. Our simulations show the mechanical unfolding process of HIV-1 TAR as an unzipping process with some minor deviation depending on solvent conditions. The picture that emerges from the all atom simulations is consistent with our earlier works.^44–46^ In accord with the experimental findings,^34^ the RNA has higher stability in *Na*^+^, thus confirming the charge density hypothesis. A systematic difference in the magnitude of the free energy is found between *Na*^+^ and *K*^+^ ions at all *f* values, as revealed in the free energy profiles (Fig. 2A-B).

**Figure 2:**
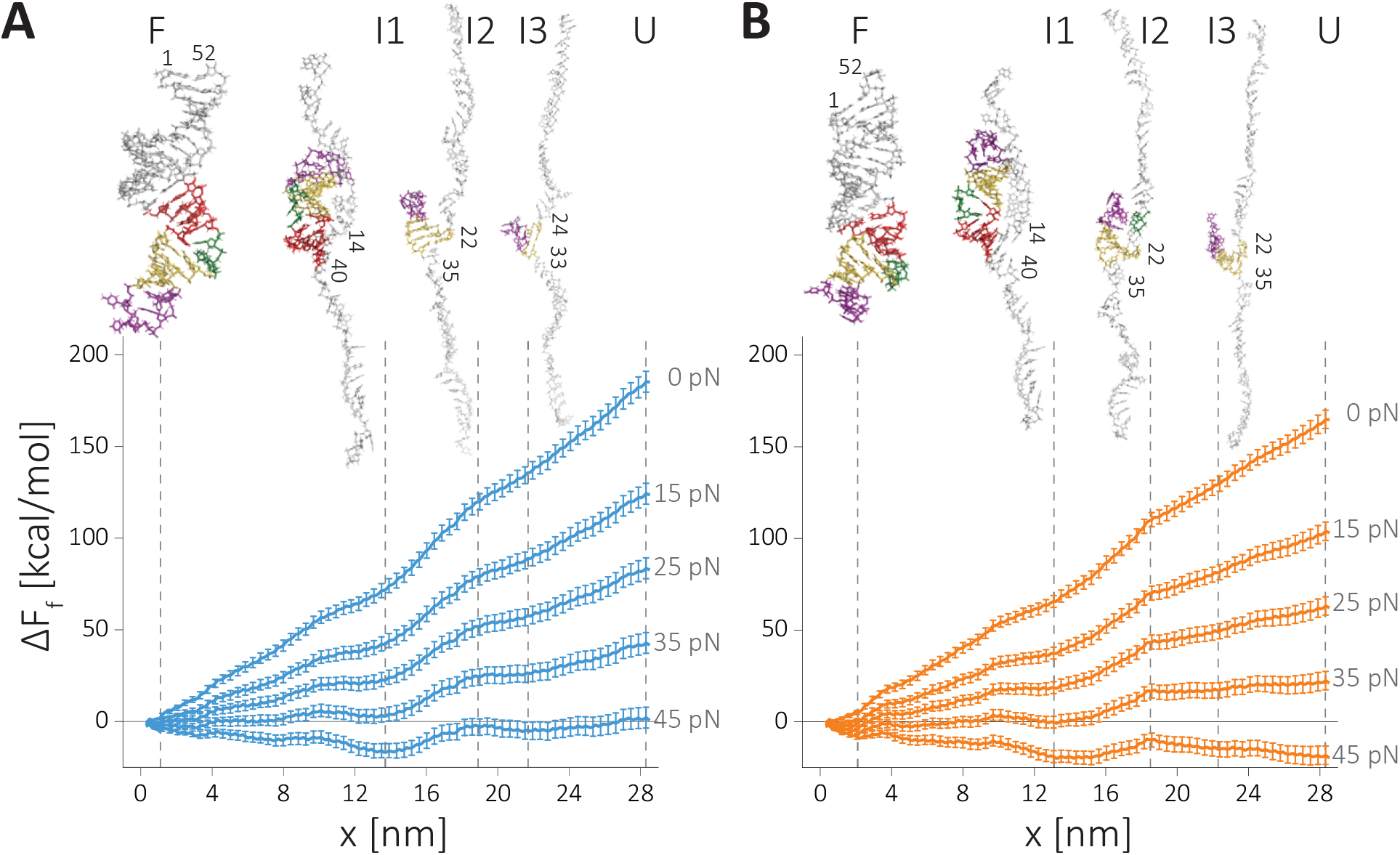
Force-induced unfolding energy landscape of HIV-1 TAR hairpin. Free energy profiles are computed by umbrella sampling MD simulations. (A) RNA in NaCl, (B) in KCl. The reaction coordinate, *x*, is the displacement along the pulling direction. The effect of force (*f*) on the landscape *F_f_* is estimated by *F_f_* (*x*) = *F*_0_(*x*) — *fx*. Vertical lines indicate major intermediate states along the RNA unfolding, F - folded state, I1 - intermediate state 1, I2 - intermediate state 2, I3 - intermediate state 3, and U - unfolded state. The snapshots from simulations representing the intermediate states are shown in the insets with the last base paired hairpin is marked.

Based on the location of the local minima, which appear *f* ≠ 0, we monitor the unfolding mechanism. The insets in Fig. 2A-B summarizes the changes in the secondary structures as RNA transitions from the folded state to the unfolded. States labeled F, I1-3 or U represent the major points along the energy landscape. The *U* is the fully extended state with no secondary structure. *I3* is partially folded state the upper stem and bulge are formed, *I2* is the putative transition state for the unfolding transition that marks the rupture of the upper stem, *I1* is the intermediate where the apical loop and lower stem are formed. In this state we found additional non-native bond formation that facilitates a higher stability of this intermediate state compared to the unfolded state (Fig. 2 and Fig. S1). In this region, additional hydrogen bonds are formed between the unzipped single-stranded and the remaining folded segment of the hairpin structure (see Fig. S2), primarily between the guanines of the single stand and the phosphate backbone of helix, and between the adenines or uracils of the single strand and the 2’-hydroxyl group of the helix (Fig. 2 Inset, Fig. S2). Finally, *F* is the fully folded state.

Transition State: Based on the computed energy landscape, we locate the transition state for the unfolding transition to be Δ*x*\* = *x_N_* – *x*\* ≈ 17.5 *nm*, where, *x_N_* and *x*\* corresponds to native state (*F*) and transition state (*I2*), respectively. Surprisingly, the value of the calculated Δ*x*\* compares favorably well with the value (≈ 17 *nm*) measured experimentally in KCl.^36^ As noted before, the simulations show that HIV-1 TAR RNA has higher stability in NaCl solution compared to KCl. The difference in the stability between the two cations, Δ*F* = *F*_*Na*^+^_ (*x*_4_) – *F*_K^+^_(*x*_4_) ≈ 9 *kcal*/*mol*, where *F*(*x*_4_) is the free energy value in *State I3* compares well with measured stability of 7 kcal/mol.^34^ Given the approximations in the simulations, and the use high loading rates, we believe that the agreement between simulations and experiments is very good.

### The determining factor in HIV-TAR RNA stability in monovalent cations

The good agreement between simulations and experiments is used as a rationale for determining the microscopic mechanism using atomically detailed simulations. Our goal is to decipher the physical principles that explain the cation size dependent stability. To test the molecular origin of the charge density hypothesis,^10,26,34^ we monitor the changes in (i) conformational entropy of RNA, (ii) the degree of RNA compaction, (iii) the energetic factors, and (iv) solvent structuring during the unfolding process. The calculation of these quantities are described in the *Computational Methods* section of the Supporting Information.

Both the conformational entropy and the degree of compaction (measured by radius of gyration) show an increase upon unfolding with a sharp transition in entropy near the transition state (Fig. S3). However, no difference is evident between *Na*^+^ and *K*^+^ ions that can explain the observed change in stability. We examined the number of water molecules released during unfolding (Fig. S4). The Radial Distribution Function (RDF) associated with water shows that hydration shell is more structured in the unfolded state. The number and binding affinity of water molecules decrease as RNA folds. However, none of the aforementioned factors show a notable cation size dependence that can distinguish between the different effects the two cations have on RNA stability.

In order to investigate the effect of the cation size, we turned our attention to RNA–ion interactions. We compute the non-bonded interaction energy as a function of the molecular extension. We also partitioned the non-bonded energy terms into electrostatic and van der Waals terms. The results, displayed in Fig. 3, show that the magnitude of the electrostatic energy far exceeds the van der Walls in both cation types, suggesting that the electrostatic interactions are dominant. Interestingly, unlike the previous analysis (Fig. S3–4), the non-bonded ion-RNA interaction energy shows marked differences between the two cations. RNA–*Na*^+^ interactions are energetically more favorable compared to RNA–*K*^+^ interactions.

**Figure 3:**
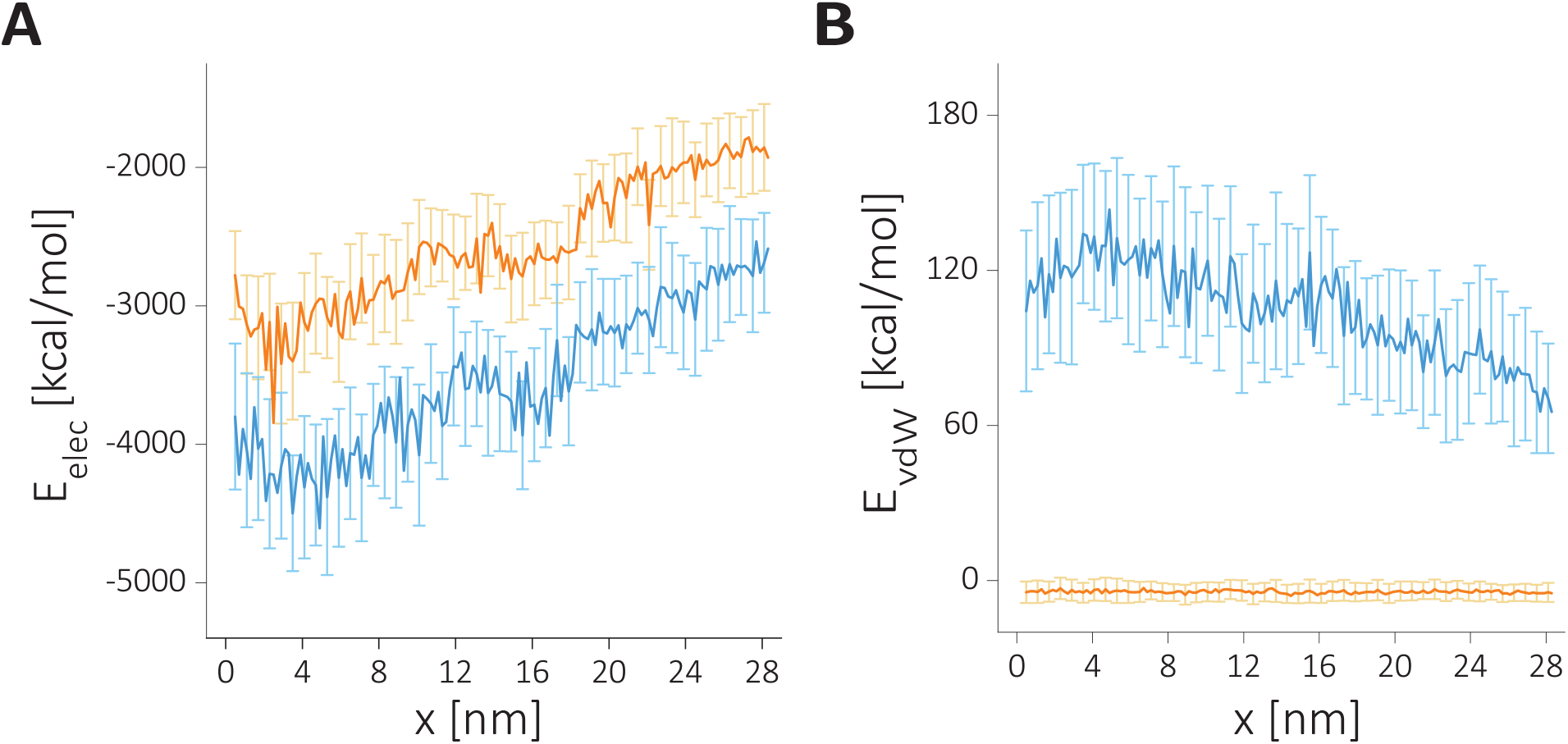
The change in the intermolecular non-bonded interaction terms as a function of extension when RNA is in *Na*^+^ (blue) or in *K*^+^ (orange). A) The electrostatic energy component. B) The van-der-Waals energy component.

### Cation size dependent ion solvation

The stark difference between the cations in the electrostatic energy offers a plausible explanation for the higher stability in NaCl. To understand the differences in the electrostatic interactions, we first investigate the cation distributions around the RNA along the mechanical unfolding pathway. Using the midpoint of the unfolding force, *f*_1/2_, we divide the RNA population into folded and unfolded states (Fig. S1). Among two minima in the free energy, we select *State I3* to represent the unfolded state. Note that our conclusions are general, and not dependent on the precise choice of a particular unfolded state. In Fig. 4A-B, we show RNA–cation distance distributions in the two basins, representing the folded and unfolded state ensembles.

**Figure 4:**
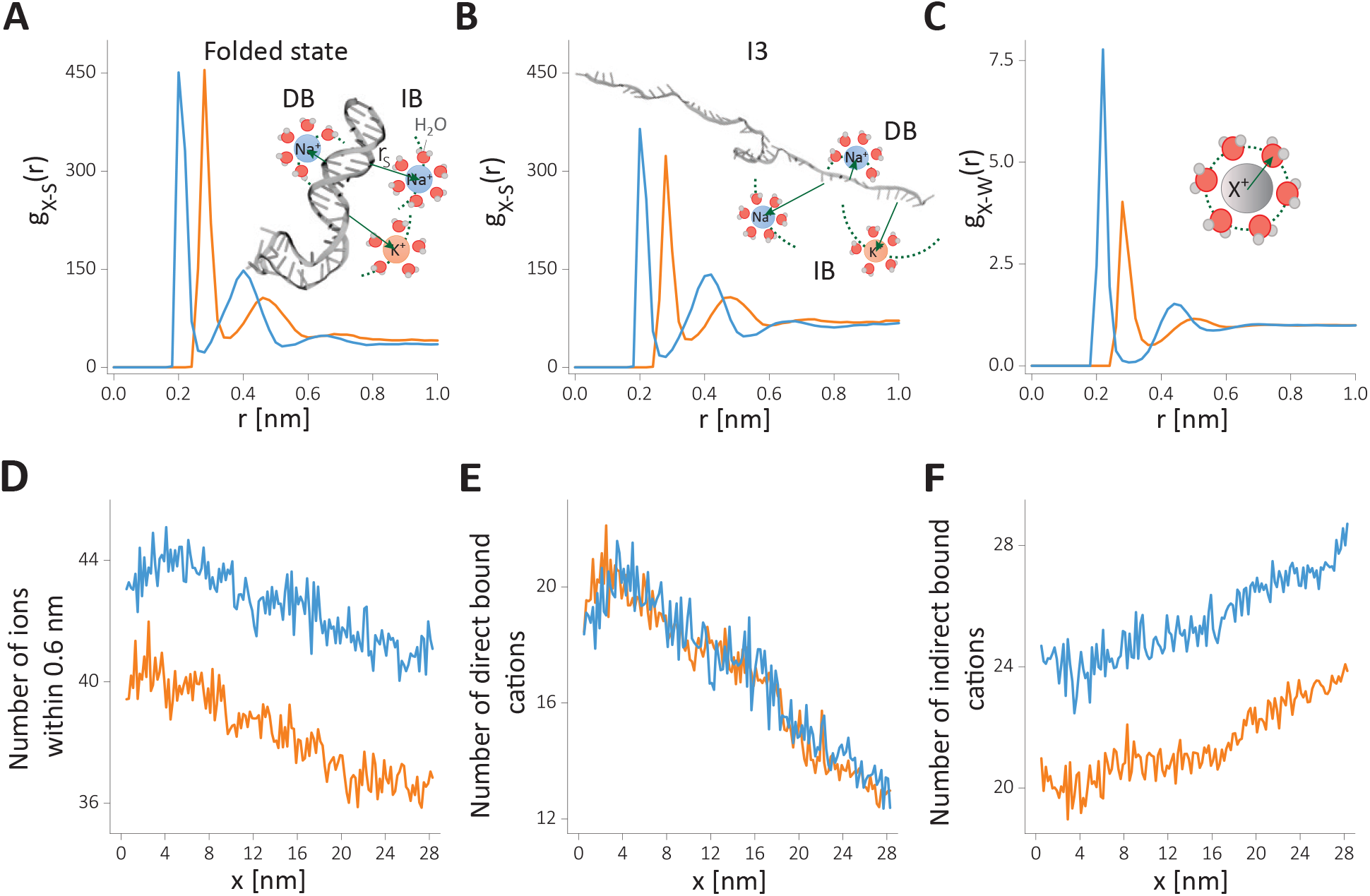
Cation distributions around the HIV-1 TAR hairpin in the folded and unfolded states. RDFs are computed from the surface of the RNA to the cation. A-B) Radial distribution function (RDF) of cation–RNA, *g_X-s_*(*r*), for, A) in the folded state and, B) in the unfolded intermediate state *I*3. Schematics in A and B illustrate the two binding modes; direct binding (DB) and indirect binding (IB). C) RDF of cations (*X*^+^) with the oxygen atom of water, *g_X–W_*(*r*). D-F) Changes in the number of *Na*^+^ and *K*^+^ coordination computed from cumulative distribution (see *Computational Methods* in the Supporting Information for details), D) the total number of ions within 0.6 nm influence radius from the RNA surface, E) the directly bound cations and, F) the number of indirectly binding cations. Data for *Na*^+^ and *K*^+^ are given in blue and orange, respectively

The RDF of cations reflects their preferential binding propensities and their binding strengths (Fig. 4A-B). We first focus on the cation distributions in the folded state (Fig. 4A). The first peak at 2.0 Å corresponds to direct binding (DB) of *Na*^+^ cations while for *K*^+^ this peak shifts further to 2.8 Å. In the DB mode, cations exchange one of the water molecules in the solvation shell with the RNA surface atom. Interestingly, despite the proximity of *Na*^+^ ion to the negatively charged RNA surface, the peak height in the RDFs is the same as in the *K*^+^ ion. Even though RNA–*Na*^+^ interactions are electrostatically favored, the smaller radius of *Na*^+^ relative to *K*^+^ also results in a stronger hydration shell. The *Na*^+^ ions disfavor dehydration of water more than *K*^+^. Hence, *Na*^+^ ions less frequently engage in direct binding (Fig. 4C) to the RNA surface. The larger size of *K*^+^ ions, on the other hand, leads to a weaker electrostatic interaction with the RNA surface. The weaker hydration shell of *K*^+^ ions also promotes direct binding of *K*^+^ ions to the RNA surface. As a result, the smaller size of *Na*^+^ does not provide an obvious advantage over *K*^+^ ions in terms of binding affinity to the first hydration shell. The second peak in Fig. 4A corresponds to the indirect binding, (IB) of cations. Here, we observe higher affinity for *Na*^+^ ions, reflecting the small hydrodynamic radius. Hydrated *K*^+^ ion that is about Δ*r* ≈0.8 Å larger in radius than *Na*^+^ ions result in a weaker electrostatic interaction, and hence a smaller amplitude in the RDF.

A similar picture emerges when we analyze the unfolded state (Fig. 4B), suggesting that the two binding modes are common in the RNA conformational states. There is a general trend of weakening of the cation binding affinities in the unfolded state, reflected in a reduction of RDF peak amplitudes. This is in accord with the reduced charge density of the backbone *ρ_c_* ∞ (1/*R_g_*) reported in Fig. S3B. Interestingly, the direct binding of *K*^+^ ions is disfavored in the unfolded state, implying a correlation between cation binding and the degree of folding. Such a correlation is not evident for *Na*^+^ ions.

The coordination number, obtained from RDFs, could be used to assess the degree of cation association. In addition, the coordination number allows us to benchmark our findings with the mean-field theories and monitor the coupling between the RNA conformations and counterion association as the RNA unfolds under tension. In Fig. 4D-F, we plot the *x*-dependent changes in the total, direct and indirect binding cation coordinations, respectively. A strong coupling between RNA conformation and cation association is evident for all the binding modes. At small *x*, corresponding to the folded state, cations accumulate near RNA (Fig. 4D). Remarkably, the number of *Na*^+^ cations show about ≈10% higher condensation. The cation size dependent difference in cation occupancy was reported in the ion competition experiments for double stranded DNA at ≈50 mM monovalent salt concentrations.^47^ Higher salt concentrations and dsRNA instead of dsDNA are expected to result in a bigger difference, which accords well with our findings. Direct binding shows a steeper change in condensation in Fig. 4E. The number of cations that are directly bound increases about two fold during the folding process, identified as a decrease in *x* while the total number of ions increase about 10%. Interestingly, the dependence of the number of bound cations to RNA as a function of *x* is independent of the cation size. In sharp contrast, the number of cations that are bound indirectly to RNA, show marked differences between *Na*^+^ and *K*^+^ ions. Our data suggests that hydrated *Na*^+^ cations accumulate more onto the RNA surface than hydrated *K*^+^ ions. The number of IB ions decreases as *x* decreases (transition to the folded from the unfolded state), suggesting a migration from the indirectly bound cation atmosphere to the directly bound state as the RNA charge density increases.

As a result of the cation size dependent condensation, *Na*^+^ ions neutralize ≈79% of the RNA charges, whereas, *K*^+^ ions shield only about ≈73% when HIV-1 TAR RNA is in the folded state. These values are in reasonable accord with the counterion condensation theory (CCT).^15^ However, CCT predictions do not readily account for cation size dependence. This implies that the subtle roles that *K*^+^ and *Na*^+^ play in affecting RNA stability requires a more nuanced picture, which we discuss below.

### Cation size dependent spatial distributions

Our analysis suggests that *Na*^+^ ions condense more onto the RNA. What are the molecular details of increased localazition of *Na*^+^ relative to *K*^+^ ions around RNA? To answer this question, we divide the RNA surface into two groups following previous studies.^48,49^ We focus on the major groove and the phosphate group of the RNA backbone. We plot in Fig. 5 the partitioning of cations into these groups.

**Figure 5:**
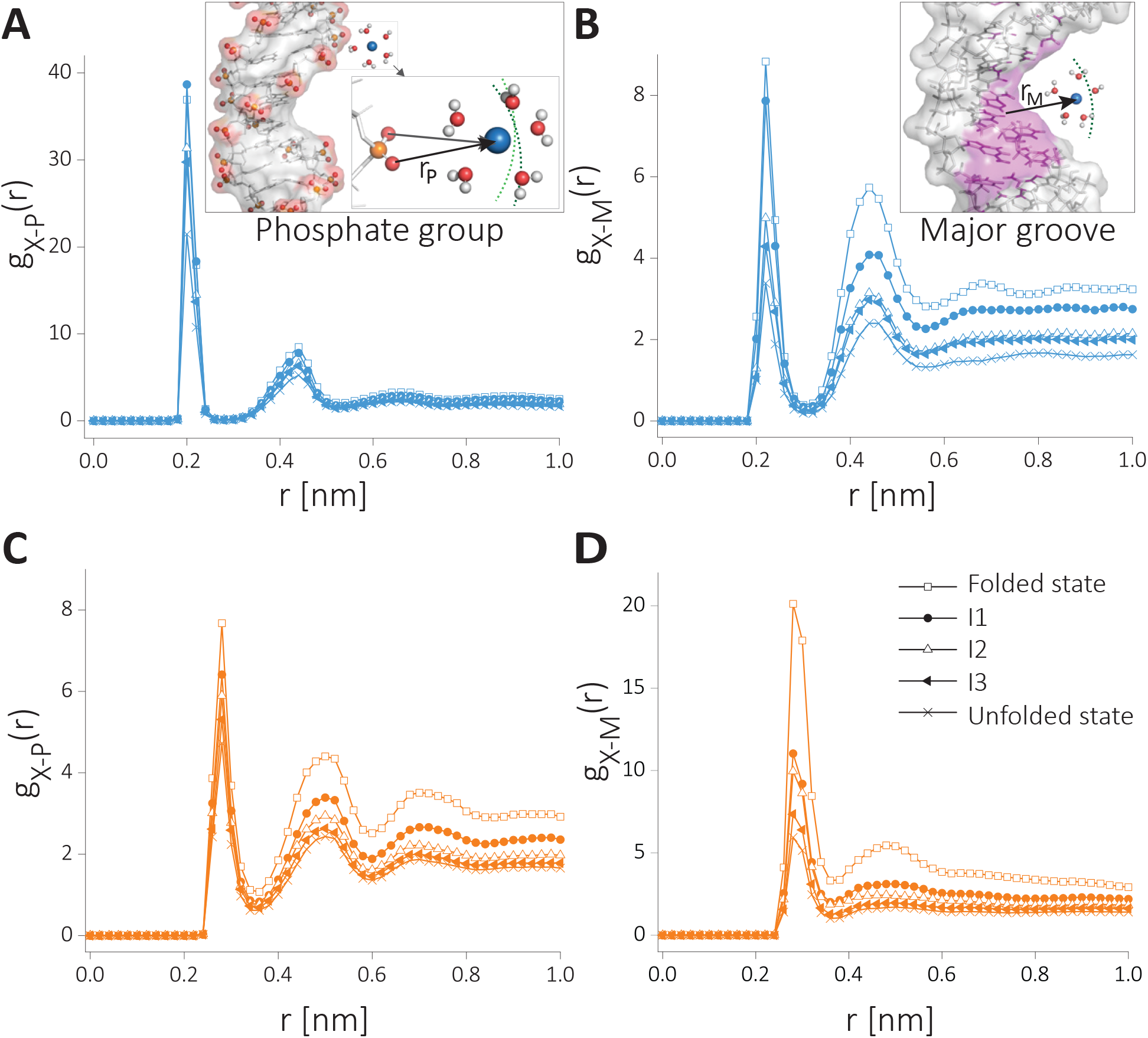
Radial Distribution Function (RDF) of cations with the two major binding sites of RNA at the defined states along the unfolding path. The RDF of A) *Na*^+^–phosphate group, B) *Na*^+^–major groove, C) *K*^+^–phosphate group, and, D) *Na*^+^–major groove.

Similar to Fig. 4A-B, the RDFs of the subgroups shows two peaks corresponding the direct and indirect binding (Fig. 5). Binding to the phosphate group is favored for *Na*^+^ ions, whereas binding to the major grooves is favored by *K*^+^ ions. When the RDFs are compared along the reaction coordinate, there is a clear correlation between the charge density of RNA backbone (Fig. S3B) and RDF peak amplitudes (Fig. 5A-B). In the case of *Na*^+^ ions, the major difference in the RDFs is in the direct binding to the phosphate groups and in the indirect binding to the major groove. In the case of *K*^+^ ions, direct binding to the major groove shows the strongest dependence on the conformations. The folded state shows a marked difference in the peak heights, suggestive of strong dependence on the RNA conformations.

### Residue level cation localization along RNA folding pathway

In Fig. 5 we show the differences in the partitioning of the cations averaged over the whole RNA surface. To gain a more detailed understanding of how the cations are localized, we zoomed-in to the residue level differences by monitoring them as RNA folds. To quantify the ionic environment, we compute local cation concentrations at the residue level (see *Computational Methods* in the Supporting Information for details). As in the previous section, we partition the binding into two subgroups. The direct and indirect binding is considered here separately. We monitor the local cation concentration as RNA undergoes structural transitions. The results for *Na*^+^ and *K*^+^ distributions are compared in Fig. 6A-D.

**Figure 6:**
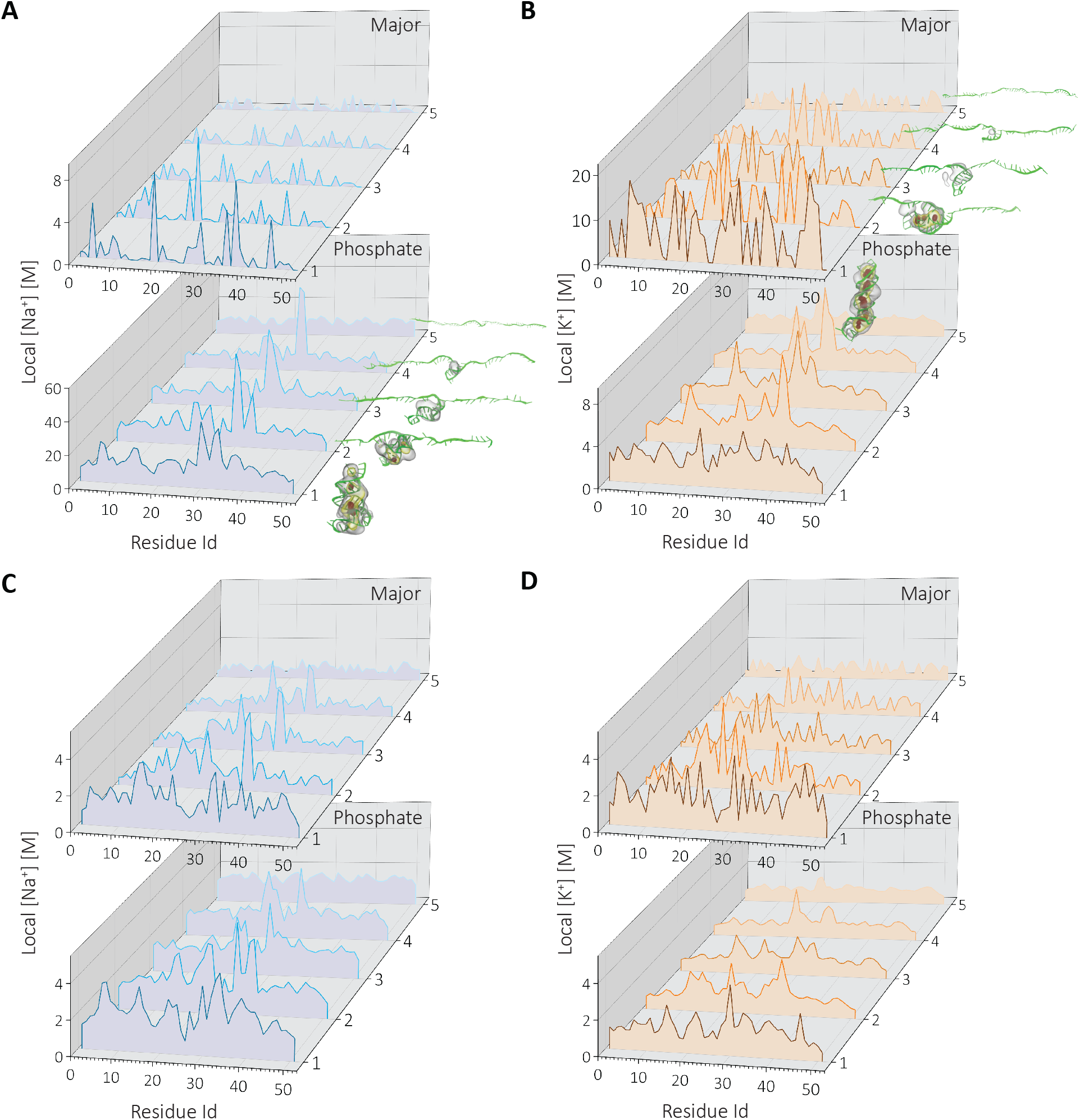
Residue level cation concentration at the defined states along the RNA unfolding pathway. The states monitored are Folded State (1), *I*1 (2), *I*2 (3), *I*3 (4) and the unfolded state (5). A-B) the concentration of direct binding cations to major and phosphate backbone, and C-D) the indirectly bound cations for *Na*^+^ (blue) and *K*^+^ (orange). Insets show local concentration in 3D density profiles at each state (see *Computational Methods* in the Supporting Information for details). Only the high-density regions are shown, *ρ* > 11_*ρbulk*_ (red), *ρ* > 7*ρ_bulk_* (yellow), and *ρ* > 3*_ρbulk_* (grey).

Although the number of direct binding cations is identical between *K*^+^ and *Na*^+^ (Fig. 4E), their partitions among the major groove/backbone shows dramatic differences. The localization of *K*^+^ ions around the phosphate group is weaker compared to binding to the major groove. In addition, *K*^+^ localization shows differences when compared with *Na*^+^ ions, despite the strong similarity between the RNA structures sampled in each cation. In accord with the results in Fig. 5, *Na*^+^ shows weak localization in the major groove in the earlier stages of folding. *Na*^+^ cation localization starts with the phosphate backbone of the bulge region (*State I3*), followed by upper and lower stem formations (*States I1-2*). Localization of the major groove, on the other hand, occurs in a later stage of folding (*State I1*), where almost all RNA is in the folded form. The high cation density regions in the concentration plots coincide well with the 3D ion densities computed for each conformational ensemble (Fig. 2, and Fig. 6 bottom panel).

In contrast, *K*^+^ ions show discrete binding to the major groove, as reflected in the high local concentrations (Fig. 6B). The cation localization is initiated in the major groove binding which is started at an earlier state (*State I3*). In KCl aqueous solution, the secondary structure formation and major groove cation localization are concomitant along the folding path. Nearly every secondary structure formation event creates a new *K*^+^ ion binding site at major grooves. The structures representing the intermediate states and 3D cation density profiles, reflecting the cation atmosphere, also highlight the unique discrete binding properties of *K*^+^ (Fig. 6B up).

In Fig. 6C-D we plot the local ion concentrations around the residues for indirectly-bound cations. Here, the concentrations are smaller compared with direct binding (Fig. 6A-B). This is due to the higher volume available to the outer-sphere cations. Similar to the direct binding mode, *Na*^+^ shows higher affinity to the phosphate groups; however, the contrast between the two binding mode is minor. Secondary structure formation enhances cation localization in the case of *Na*^+^. The difference between the two cations lies the way at which they associate with the phosphate groups. *Na*^+^ shows a structure-dependent discrete binding, while *K*^+^ ions form a uniform distribution surrounding the residues at early stages of folding. However, the discrete nature of the major groove binding is still evident throughout the folding process in the case of *K*^+^.

### Sequence effects on cation localization

To quantify the sequence dependence of cation binding, we calculated local cation concentration for each type of nucleobases (Fig. 7). We observe a structure-dependent relationship for sequence dependence. Phosphate binding shows weak specificity both in the folded and unfolded states. *Na*^+^ ions interact stronger with phosphates and binding gets stronger as phosphate charge density increase at the folded state. *K*^+^ ions on the other hand do not show any significant binding when the RNA is in the unfolded state. In contrast, binding of *K*^+^ ions to the major groove shows stronger dependence at the folded state showing sequence specificity, favoring G, U, and A nucleotides.

**Figure 7:**
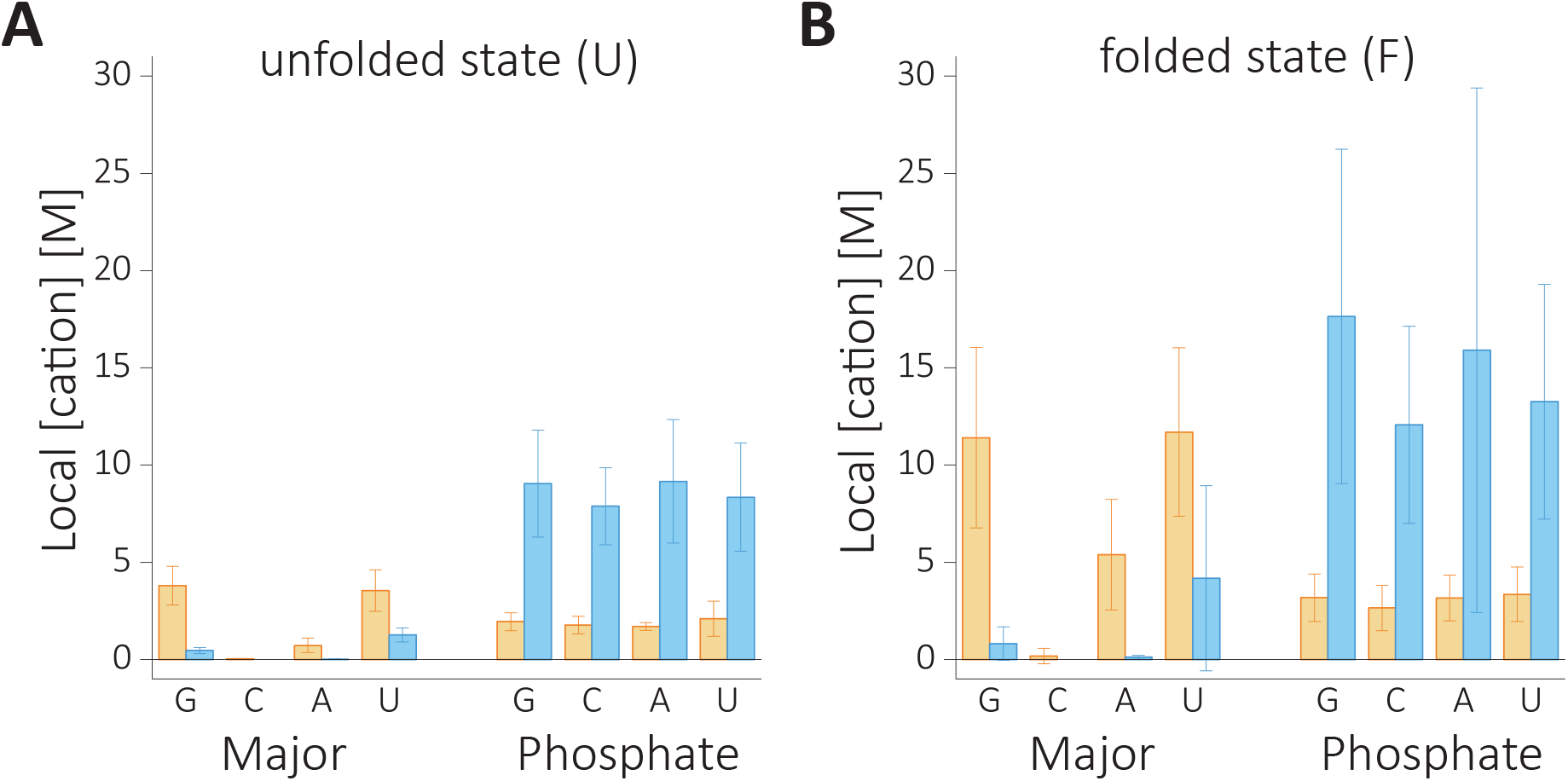
Local cation concentrations of *Na*^+^ (blue) and *K*^+^ (orange) as a function of sequence at the unfolded and folded states. Shown in A) is the unfolded state, and B) is for the folded state.

### Cation dependent accessible surface area along the RNA

By using all-atom simulations, we found a clear difference in the stability of HIV-1 TAR hairpin in KCl versus NaCl solutions (Fig. 2), which accords well with the experimental findings.^34^ After having identified that ion size dependent electrostatic interactions are the main cause for the RNA structure stabilization, we dissected the molecular mechanism leading to the observed stability in *Na*^+^ compared to *K*^+^. The major finding that explains the stability difference is that the smaller size *Na*^+^ with higher charge density ions bind preferably to the backbone phosphates, whereas larger size *K*^+^ ions with a smaller charge density prefer the major grooves. The first is an electrostatic phenomenon while the latter is a size or excluded volume effect. While *Na*^+^ condensation depends on RNA charge density, *K*^+^ ion association depends upon the secondary structure and sequence identity.

The difference between the number of bound *Na*^+^ and *K*^+^ ions to RNA is due to the stronger interactions involving hydrated *Na*^+^ ions, as well as the bigger surface area that these ions can access. To demonstrate that this is the case, we compute the average surface accessible area (SASA) available for each subgroup. The results, summarized in Fig. S5, show that *Na*^+^ ions that bind to phosphate groups have access to an area of 76 nm/s2/N in the unfolded state. This number reduces only by about 3% when RNA folds. Although the SASA remains similar, the RNA charge density increases due to compaction. In contrast, the SASA for *K*^+^ ions that primarily bind to the major groove is 55 nm/s2/N in the unfolded state. This number shows a dramatic reduction (~ 35%) upon compaction. In the folded state the SASA becomes 37 nm/s2/N, which is nearly half of the SASA available to *Na*^+^ ions. As a result of preferential binding that is dictated by the cation size, *Na*^+^ ions has larger entropy by exploring a bigger area of binding compared to *K*^+^ ions. Moreover, this binding mode cannot compensate for the weaker (compared to *Na*^+^) interaction with RNA.

## Conclusion

Taken together, we propose a novel molecular mechanism for explaining the enhanced stability of the HIV-1 TAR RNA in *Na*^+^ compared to *K*^+^. The *Na*^+^ ions preferentially bind to the phosphate groups whereas the *K*^+^ ions are localized near the major grooves. The very striking changes in the binding modes between the ions to RNA is sufficiently different, which readily explains the observed stability differences. There are two additional predictions that arise from our study. (1) Although the study focused on HIV-1 TAR, the mechanism should hold for other RNA molecules as well DNA because it depends on generic properties of cations and the structures of nucleic acids, that are stabilized by monovalent cations. (2) Because charge density of the cations determine the binding modes to RNA, we predict that the stability of RNA should decrease as one goes down the alkali metals in group I of the periodic table. RNA stability should be maximum in *Na*^+^ solution and decrease as the radius of the alkali metal ions increases.

## Acknowledgement

This work was supported by NYUAD Faculty support grant to SK. This research was also conducted with support from the NYUAD Center for Advanced Computing. DT acknowledges support from the National Science Foundation (CHe-1900033) and the Collie-Welch Chain (F-0019).

## Supporting Information

### Computational Methods

#### Molecular Dynamics (MD) simulations

We model the starting structure of the 52-nucleotide long HIV-1 TAR RNA, with the sequence 5’GGCUCUGGUUAGACCAGAUCUGAGCCUGGGAGCUCUCUGGCUAACU AGGGCC-3’, by extending the crystal structure of HIV-1 TAR RNA (PDB entry: 1ANR; residue 19-43).^50^ The double-stranded (ds) RNA construct comprising the sequences of 5’-GGCUCUGGUUAGAC-3’ and 5’-GCUAACUAGGGCC-3’ were built using the Nucleic Acid Builder.^51^ We joined the two units manually and minimized the energy to refine the structure. For this and all other calculations, we used GROMACS 5.05 program with ff99-parmbsc0 forcefield^52^ due to its success in capturing the HIV-1 TAR dynamics in the previous study.^53^

The energy-minimized HIV-1 TAR hairpin structure was aligned along the *x*-axis of a periodic box with dimensions 42×6.5×12.5 *nm*^3^. The dimension along the long axis is chosen to ensure that the RNA does not interact with the periodic images after it is fully stretched, which is technically important in performing the pulling simulations.

We added explicit water and ions to neutralize the RNA charge, and to mimic the experimental concentration of 400 mM monovalent salt (Fig. 1C). We created two systems, one RNA in NaCl aqueous solution and the other is in KCl with the same ionic strength. We kept the number of atoms constant in each simulation to allow direct comparison of the extensive properties of RNA in *Na*^+^ and *K*^+^. We added 771 *Cl*^−^ and 822 (*Na*^+^/*K*^+^) to the simulation boxes by replacing some of the water molecules. We used ff99paramsbc0 force field to represent the RNA,^52^ Smith and Dang parameters^54^ for ions, and TIP3P^55^ for water.

The energy of the solvated systems were minimized for about 5000 MD steps using the steepest descent method. This process eliminated high energy contacts that may arise due to random placement of water and ions. The minimized structures were then equilibrated as explained below.

First, we employed a 2.5-ns long simulation in isothermal – isobaric ensemble (NPT) by keeping the temperature at 300 K using the Berendsen thermostat.^56^ Parrinello-Rahman barostat^57^ was used to maintain the pressure at 1 bar. The heavy atoms of the RNA were restrained to their initial positions using harmonic restraints with a force constant of 1000 kJ/*nm*^2^ while ions and water were allowed to move freely. Periodic boundary conditions were implemented in the three directions. Particle Mesh Ewald (PME) summation^58^ was used to compute long-range electrostatic interactions. The real space distance cutoff (for electrostatics and van der Waals energies) was set to 11Å. The grid for the Fourier space summation in the PME was 1.6^Å^, and fourth order splines were used to interpolate the charge density on the grid. A dispersion correction was made for the van der Waals cutoff. Covalent bonds in the water and RNA were constrained to their equilibrium geometries using SETTLE^59^ and LINCS^60^ algorithms, respectively. The equations of motion were integrated using the Leap-Frog scheme with a time step of 1 fs.

The dimensions and the positions of atoms at the last snapshot of the NPT simulation were saved and used to initiate the solvent equilibration step. This procedure ensures that ions and water molecules reach equilibrium before the production runs. For this purpose, we used a further restrained, 60-ns long NVT run keeping all the settings from the previous section the same except the barostats were turned, off and Velocity scaling was employed.^61^

#### Steered MD simulations (SMD)

The last frame in the equilibrated trajectory for each cation condition was used to generate the unfolding pathway of the HIV-1 TAR upon application of the mechanical force. In particular, a 300 ns long steered MD simulations^62^ were performed to pull the 3’ and 5’ termini of the RNA (Fig. 1C). For the reaction coordinate we used the molecular extension, which is the *x*-component of the center of mass distance between the two termini nucleotides. To realize as slow a pulling speed as possible in the MD simulations, we used a pulling rate of 0.1 nm/ns with a force constant of 1000 kJ/(mol×*nm*^2^).

#### Free energy calculations

Using the conformations along the force-induced unfolding pathway of the SMD we construct the free energy profile of RNA as a function of the pulling coordinate, *x* using umbrella sampling MD simulations.^43^ Each conformation picked was restrained using a harmonic biasing potential *U* = (*k*/2)(*x* — *x_i_*)^2^, where *k* =1000 kJ/(mol×*nm*^2^), and *x_i_* is the reference distance of the *i^th^* window. We used about 142 windows with a 2Å inter-window spacing, and we sampled conformations for about 100 ns in each umbrella, giving rise to an aggregate simulation time of ≈14 microseconds for each solvent condition. The sampled conformations were saved every 10 ps for data analyses.

The free energy landscape was constructed by calculating the potential of mean force (PMF) using the GROMACS-implemented Weighted Histogram Analysis Method (WHAM).^63,64^ Statistical errors were estimated using the bootstrap analysis with 200 bootstrap samples. From the PMF, we estimated the force dependent free energy change along *x* at various applied forces.

#### Intermolecular interaction energy

We used a NAMDEnergy analysis tool (https://www.ks.uiuc.edu) implemented in VMD to calculate the non-bonded intermolecular interaction energy for each umbrella window. Coordinates of the atoms were extracted every 1,000 ps. The data in each umbrella is divided to equal pieces to estimate the standard error presented as mean±S.D.

#### Surface bound water and ion numbers

Radial distribution function (RDF) of group X around group Y, *g_X-Y_*(*r*), was calculated to gain insights into ion and water distributions. The cumulative number of cations on the RNA surface (S) up to distance *R*, *N_X-S_*(*R*), is computed as:

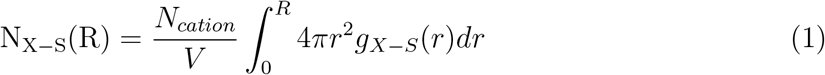

where *N_cation_* is the total number of cations in the simulation box volume *V*, *R* is the cut-off distance beyond which the cation is deemed to be unbound. To find the number of inner-sphere cations (NIS) we chose *R* to be the minimum between the first two peaks in the RDFs. Note that this number depends on the cation size. For outer-sphere cations, we integrate the equation to the second minimum and subtract the value the from NIS. Similarly, water molecules around RNA surface (S-W) or cation (X-W) were investigated. All atoms of water were considered for water-RNA interactions, whereas for the investigation of cation-water, we used oxygen atom of the water.

#### Configurational entropy

We applied principal component analysis (PCA) to elucidate collective motions from simulation. All heavy atoms of the RNA were used in performing the PCA. We diagonalized the covariance matrix of atomic fluctuations along the folding pathway. The RNA chain entropy, *S*, is estimated by analysing the resulting eigenvalues, *σ*, using the Schlitter’s formula^65^:

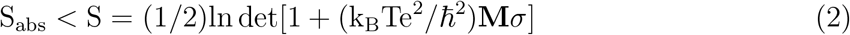

where *k_B_* is the Boltzmann’s constant, *ℏ* is the Planck’s constant, and *T* is the absolute temperature, *e* the Euler value, and **M** is the diagonal mass matrix of rank 3*N*.

#### Local ion concentration profiles

To provide further details on cation localization around RNA we compute local cation concentration. For that, we divided RNA into two subgroups: cations around the phosphate backbone, and cations around major grooves. The concentration around each group is estimated as,^66^

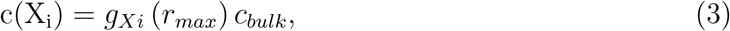

where, *X_i_*≡{Phosphate group, Major groove}represents the two subgroups for residue *i*. We used O1P and O2P atoms to represents the backbone phosphate group. The major groove atoms are nucleotide dependent. We chose for A(N6, N7), C(N4), G(N7, O6), U(O4). *gx_i_*(*r_max_*) is the maximum peak height of the normalized RDF of the cation and atom(s) on the RNA subgroup, *c_bulk_* is the bulk solution concentration.

#### Cation density profiles

To visualize the cation cloud around the RNA, we computed the cation density in Cartesian coordinates. For an arbitrary point in space **r** ≡ (*x*, *y*, *z*), the average cation number density *φ*(*x*,*y*,*z*) is computed from the trajectory using,

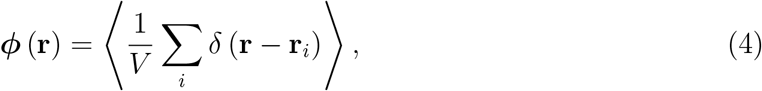

where the sum goes over the cation index *i* in the simulation box of volume *V*. *δ*(*x*) is the Kronecker delta, and 〈…〉 represents the ensemble average.

### Supplementary Figures and Analysis

**Figure S1:**
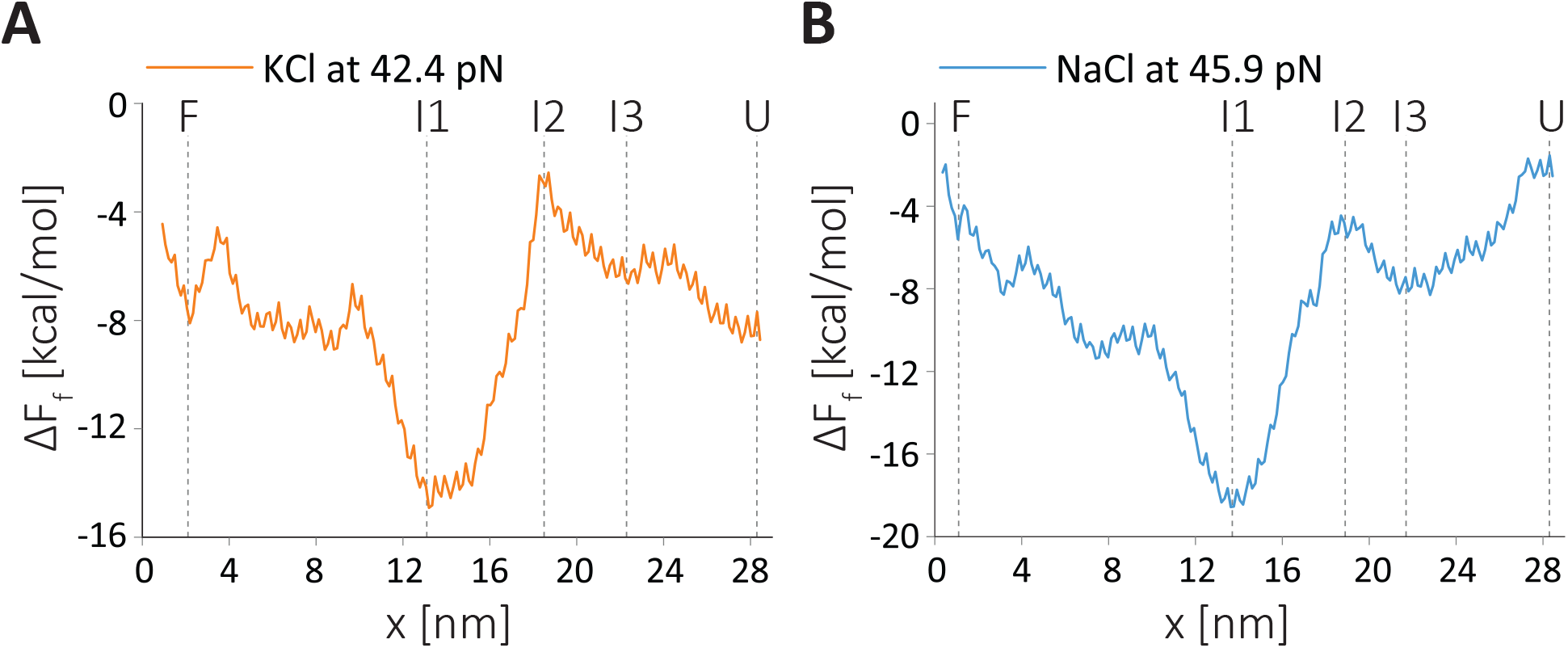
Free energy at the unfolding transition where folded and unfolded populations of RNA are equal. The effect of force (*f*) on the landscape *F_f_* is given as *F_f_*(*x*) = *F*_0_(*x*) — *fx*, with *x* the displacement along the pulling direction. (A) is for *K*^+^, and (B) is for *Na*^+^. Major intermediate states are represented by vertical lines that explained in Fig. 2. The minimum at the midpoint force corresponds to *I*1 state where additional stabilization occurs due to non-native contact formation explained in Figure S2.

**Figure S2:**
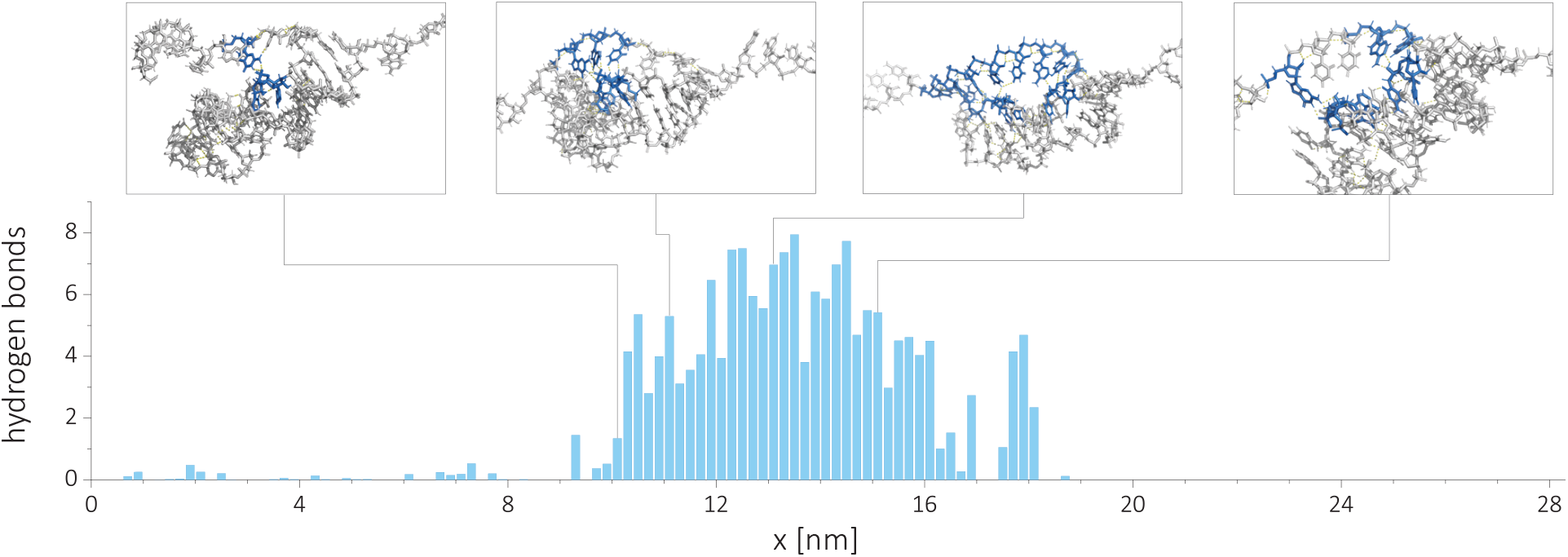
Average number of inter-residue hydrogen bond formation as a function of extension for HIV-1 TAR hairpin in NaCl. The average number of hydrogen bonds were computed between segments 6U-14C and 21U-37C. Insets show representative structures at *x* ≈10.1, 11.1, 13.1, and 15.1 nm. The residues that form hydrogen bonds are highlighted in blue. Similar results obtained for RNA in KCl

#### Conformational entropy and radius of gyration analysis

To elucidate the mechanism of cation size on RNA stability we compute the conformational entropy and the degree of compaction as a function of extension. As the RNA unfolds the conformational entropy increases (Fig S3A). A linear increase in the entropy between *x* ≈ (0 – 9*nm*) coincides with the unzipping of the extended stem region. Interestingly, unzipping in the range *x* ≈ (9 — 16 *nm*) results in a smaller change in entropy because it involves disruption of 3-nt apical loop region, which is structurally already disordered even in the folded state. The rupture of the base pair 36U-18A (Inset in Fig 2A-B) results in an abrupt increase in entropy. The rupture coincides well with the transition state location (Fig. 2A-B). After the transition state, the entropy change shows a linear increase with extension.

The change in the radius of gyration (*R_g_*) of RNA on the other hand, is impervious to increase in the extension at *x* ≈ (0 — 8 *nm*), as is evident from the nearly flat region in Fig. S3(B). After the rupture of the lower stem (Fig 2A-B) the *R_g_* ramps up. However, neither entropy, nor radius of gyration show cation size related differences.

**Figure S3:**
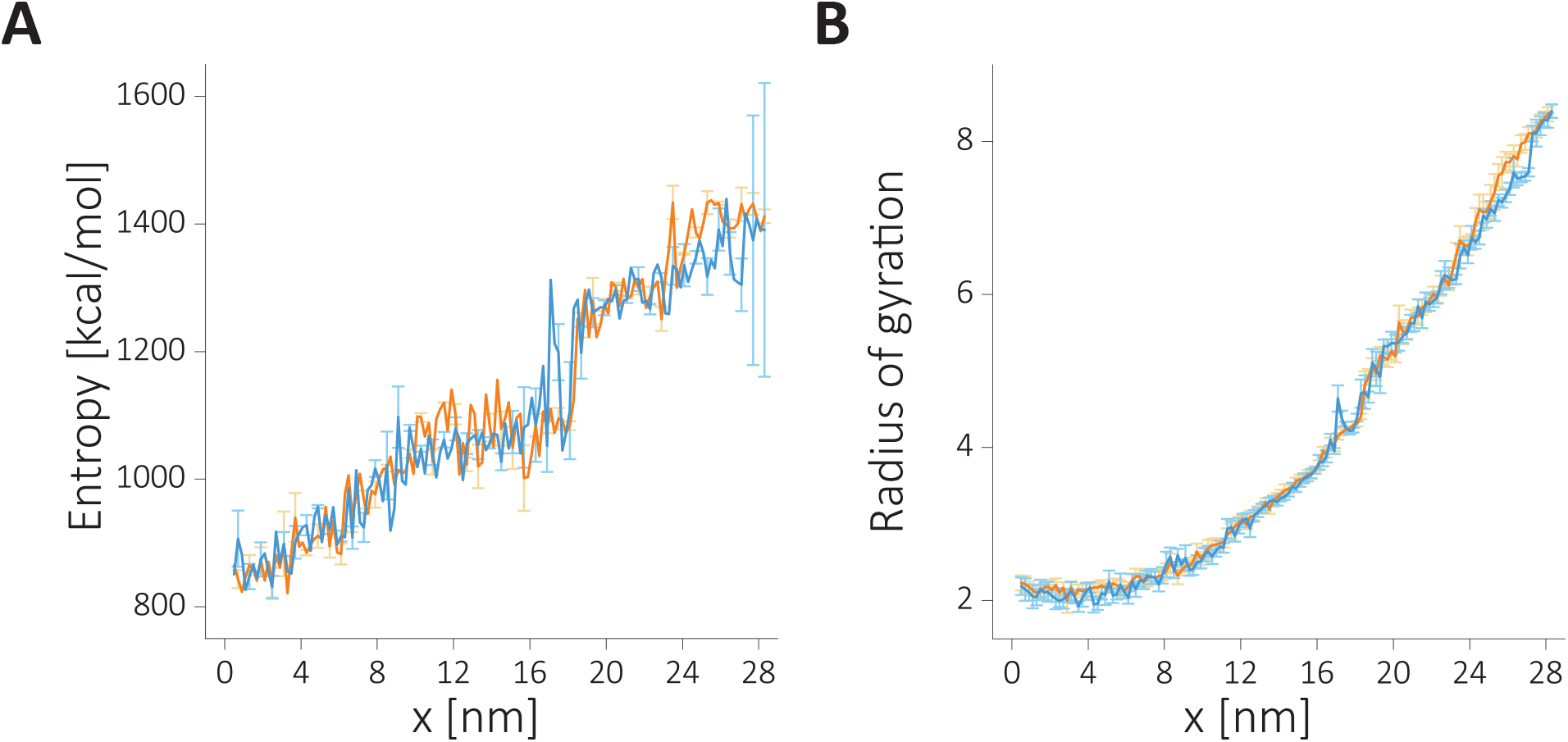
The change in the RNA conformational entropy and RNA compaction as RNA undergoes unfolding transition with applied force. The RNA in the presence of KCl (orange) or in NaCl (blue).A) The change in the RNA conformational entropy in (*TS*) as a function of extension. B) The change in the radius of gyration during the unfolding process.

**Figure S4:**
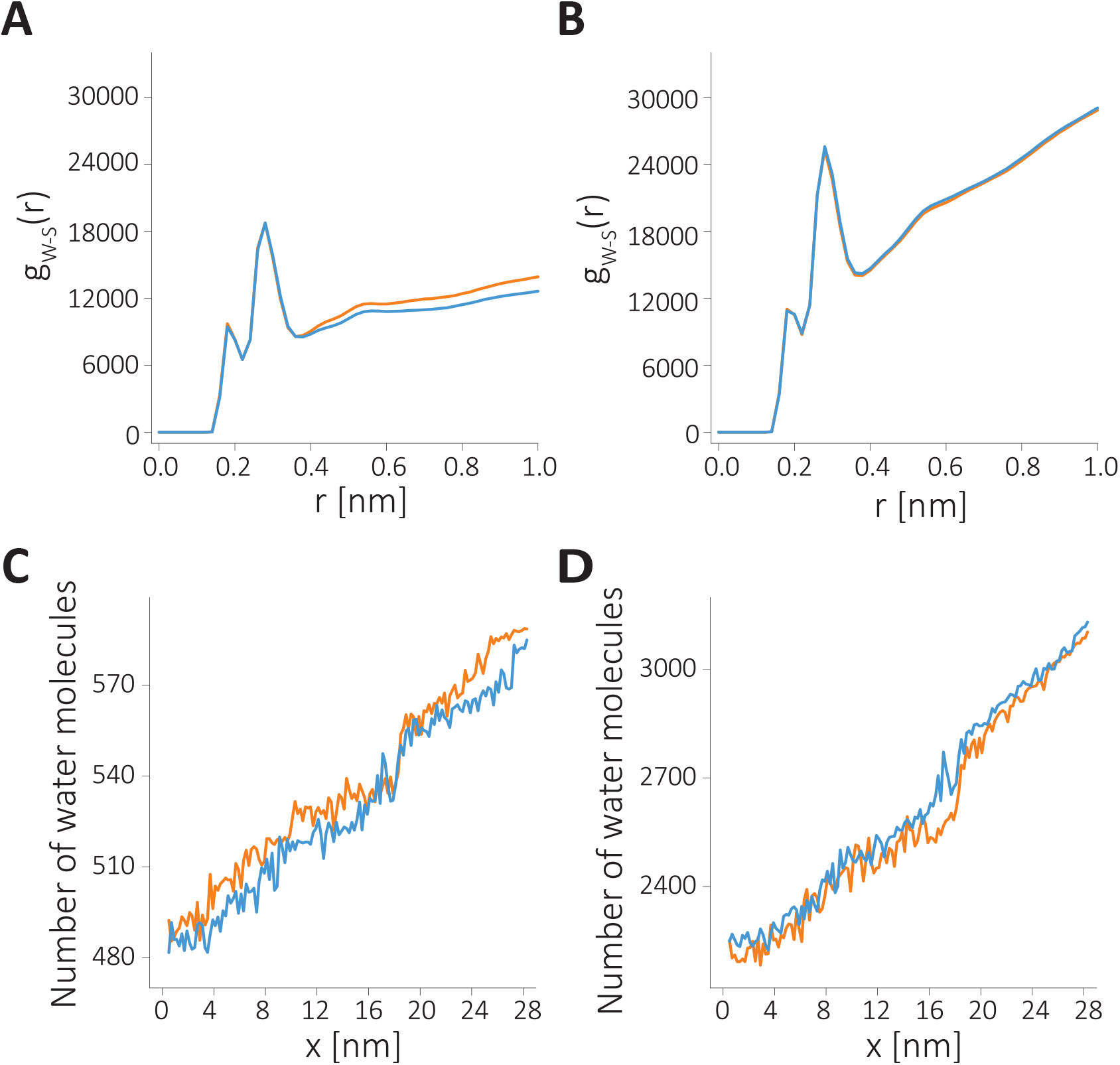
Radial distribution function (RDF) of water molecules around the surface of HIV-1 TAR hairpin at the folded and unfolded states. Data for *Na*^+^ and *K*^+^ are given in blue and orange, respectively. A-B) RDF of water (W) with the RNA surface (s) at the folded and at the unfolded states respectively. (C-D) Changes in the cumulative number of water molecules around RNA as a function of extension. The number of surface-bound water molecules was calculated based on equation 1. C) Directly binding water coordination (set by cut-off: 0.22 nm) and, D) in directly binding water coordination (cut-off: 0.36 nm).

**Figure S5:**
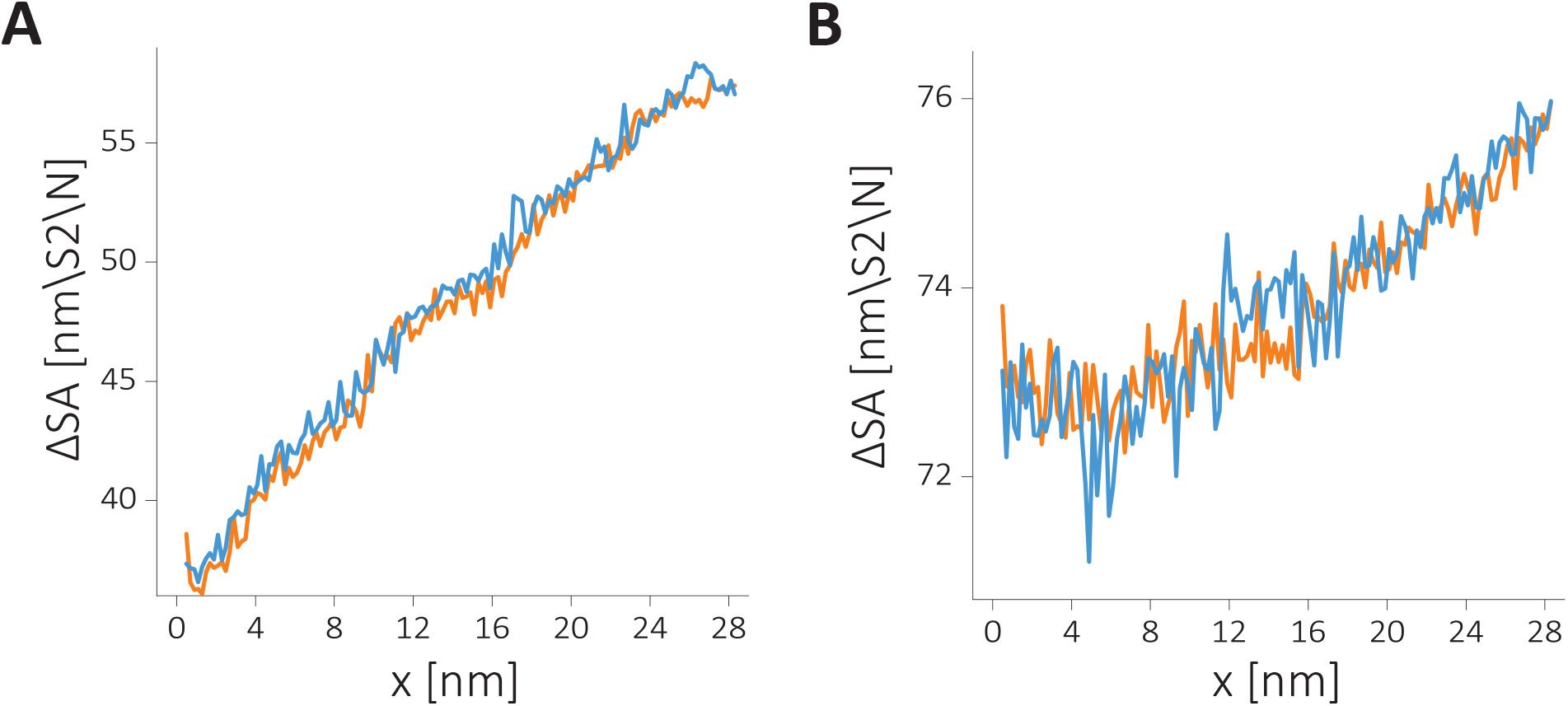
Solvent accessible surface area (SASA) computed along the unfolding pathway. A) SASA of the major grooves and, B) SASA of the phosphate groups. *Na*^+^ (blue) and *K*^+^ (orange)

## Graphical TOC Entry

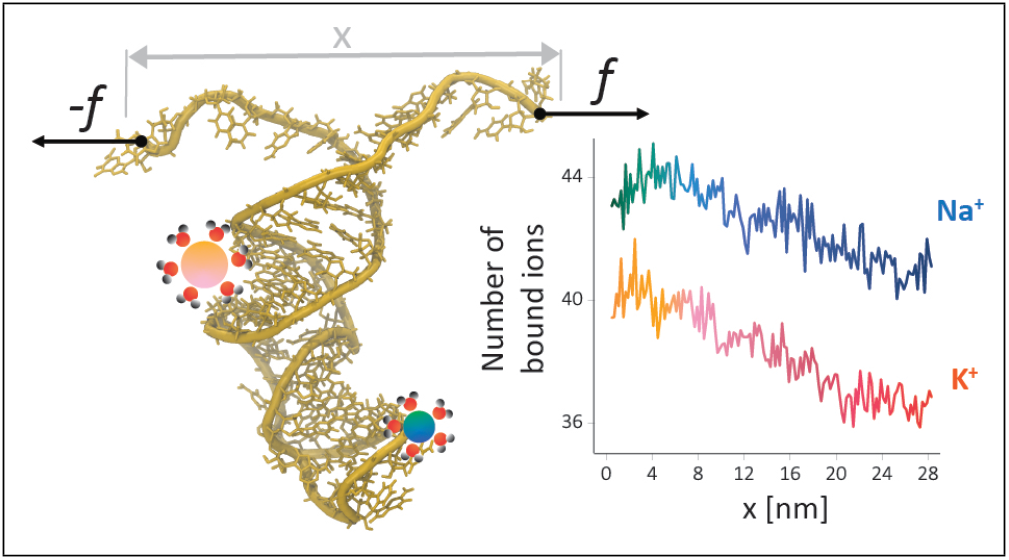

